# samsampleX: Distribution-aware downsampling for benchmarking next-generation sequencing data

**DOI:** 10.64898/2026.06.03.729942

**Authors:** Sedat Demiriz, Daniel Taliun

## Abstract

**Summary:** High-throughput next-generation sequencing (NGS) is essential for genetic variant discovery across diverse applications. As NGS evolve, there is a growing need for benchmarking tools that support realistic data simulation and downsampling. Existing downsampling tools apply uniform sampling of sequencing reads, which inadequately models realistic coverage distributions, particularly in difficult-to-sequence regions and hybrid sequencing designs. Here we present samsampleX, a Python-based tool implementing a novel distribution-aware downsampling algorithm that dynamically adjusts read retention probabilities to emulate coverage profiles derived from real sequencing data. Using ultra-high-coverage reference datasets, samsampleX accurately reproduces coverage patterns observed in typical sequencing experiments, outperforming uniform downsampling methods at preserving depth variability across genomic regions such as the HLA locus and hybrid whole-exome/genome sequencing configurations. samsampleX extends current downsampling strategies by offering enhanced flexibility for specialized NGS benchmarking scenarios, facilitating improved assessment of sequencing data analysis methods.

**Availability and Implementation:** samsampleX source code, benchmarks and usage instructions are available at https://github.com/sdemiriz/samsampleX under an MIT License.

## 1. Introduction

High-throughput Next Generation Sequencing (NGS) technology has become foundational for genetic variant discovery and assessment across a wide range of applications, including genome-wide association studies (Yi et al. 2024; Canela-Xandri et al. 2018) and molecular diagnoses of rare diseases (Satam et al. 2023; Karczewski et al. 2020). As new sequencing techniques are developed to reduce costs and improve the precision of variant discovery (Bhérer et al. 2024; DeFelice et al. 2024), their adoption depends on the availability of standardized datasets (Mangul et al. 2019) and compatible tools that enable thorough benchmarking (Wagner et al. 2022).

A central parameter of NGS data is depth of coverage (DP), defined as the average number of times each DNA base-pair is sequenced. For instance, high-DP whole-genome sequencing (WGS) assesses each DNA base-pair around 33 times (i.e. DP=33×) and allows the detection of single-nucleotide variants with >99% accuracy (Rashkin et al. 2017). Higher DP further enhances confidence in clinical applications (Petrackova et al. 2019). Population-based genetic studies may sequence thousands of individuals at lower DP to reduce sequencing costs, accepting a 20-60% loss in variant detection power (Baraja-Fonseca et al. 2025), which is compensated for by statistical imputation of missing genotypes (Pasaniuc et al. 2012). Whole-exome sequencing (WES) reduces sequencing costs by restricting coverage to protein-coding regions, which comprise approximately 1% of the genome, thereby excluding non-coding variants when they are not relevant to the study’s hypothesis. Emerging hybrid methods, such as Whole Exome Genome Sequencing (WEGS) (Bhérer et al. 2024) and Blended Genome Exome (BGE) (DeFelice et al. 2024), combine low-DP WGS with high-DP WES to provide a cost-effective middle-ground alternative.

Gold-standard NGS datasets, such as Genome In A Bottle (GIAB), include harmonized data at ultra-high-DP, ranging from 300× for WGS to 1000× for WES (Zook et al. 2016), allowing evaluation of sequencing technologies and parameters early in genetic study design. They are used as truth datasets for benchmarking *in vitro* experiments (Wong et al. 2025) or as high-DP input datasets for *in silico* downsampling (Harvey et al. 2023; Sun et al. 2021; Yu et al. 2023; Regier et al. 2018) to emulate and benchmark datasets across a range of DP levels, providing substantial cost and time savings compared to repeated *de novo* sequencing. However, popular software tools for *in silico* downsampling of aligned sequencing reads, such as GATK Picard (McKenna et al. 2010), samtools (Li et al. 2009), and sambamba (Tarasov et al. 2015), are limited to applying a single global downsampling coefficient uniformly across the input dataset (e.g. ultra-high-DP WGS). Although this approach is valid in many scenarios, it results in a DP distribution that does not reflect the variability observed in true *de novo* sequencing at lower DP, especially when using highly optimized gold-standard high-DP datasets. This may lead to biases when downsampling challenging-to-sequence regions and emulating novel hybrid sequencing technologies in existing gold-standard datasets.

In this work, we introduce a new downsampling algorithm that allows for non-uniform, pattern-aware downsampling of sequencing data. We showcase scenarios where our algorithm is more suitable than existing uniform downsampling methods. We also provide the implementation of our algorithm through a freely accessible Python-based tool, samsampleX, which offers runtime and memory use comparable to the most popular uniform downsampling tools.

## 2. Approach

The uniform downsampling algorithms implemented in GATK Picard, samtools and sambamba accept a downsampling coefficient, a value between 0 and 1, that specifies the fraction of sequencing reads from the input NGS dataset to retain. As they stream reads, these tools compute a hash value from each read name: MurmurHash3 (Appleby 2015) in GATK Picard, a combination of times-31 (X31) and Wang hashes (Wang 2007) in samtools, and SuperFastHash (Hsieh 2008) in sambamba. The resulting integer is then conceptually divided by a maximal hash value (e.g. 2^32^ in GATK Picard) to produce a deterministic pseudo-random value between 0 and 1 for each read. This value serves as the read’s probability of inclusion. Reads with hash-derived values below the specified downsampling coefficient are retained, resulting in uniform and reproducible downsampling. However, because this is a stochastic process, the actual proportion of retained reads may vary slightly between runs. To improve downsampling accuracy, GATK Picard implements two additional downsampling strategies, termed High Accuracy and Chained, alongside the standard approach (ConstantMemory in GATK Picard). The two strategies improve downsampling precision by buffering or oversampling reads at the cost of increased memory use.

In our new algorithm, implemented in samsampleX, we propose a new strategy for hash-based downsampling of sequencing reads that extends uniform downsampling and enables the creation of arbitrary DP patterns in a downsampled dataset. Instead of a single downsampling coefficient, the algorithm accepts a target DP value *d*_*i*_ for each genomic position *i* = 1, …, *N*. For each position *i*, downsampling coefficient is defined dynamically as *k*_*i*_ = *d*_*i*_/*c*_*i*_, where *c*_*i*_ ≥ *d*_*i*_ is the observed DP value at position *i* in the input dataset. *k*_*i*_ represents the fraction of reads from the input dataset at position *i* that must be retained to achieve the target DP *d*_*i*_ in the downsampled dataset and can also be interpreted as the probability of retaining a read at that position. For a read of length *L* starting at position *j*, its per-read downsampling coefficient is defined as 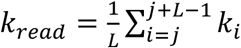, which corresponds to the expected probability of retaining the read at any position it spans. As in existing tools, our algorithm computes a hash-derived value (specifically, xxHash (Collet, 2014)), representing each read’s probability of inclusion, which is then compared to the per-read downsampling coefficient *k*_*read*_ to determine whether the read should be kept in downsampling dataset. Using *k*_*read*_ as a threshold ensures that the expected depth of coverage *d*_*i*_ at position *i* is preserved in the downsampled dataset, assuming that all reads in the input dataset are of similar length (i.e., all long or all short reads) and that *k*_*i*_ along positions of a single reads are relatively similar, which is common for short reads, but potentially violated for long reads (see Supplementary Information).

Users provide the desired target DP values, *d*_*i*_, in the Browser Extensible Data (BED, Kent et al. 2002) file format. In samsampleX, we provide functions to generate BED files from existing sequencing data, with optional smoothing by aggregating similar target DP values across nearby positions. Downsampling while maintaining an even DP is supported by providing a single desired DP value for the target region (i.e., conceptually, samsampleX is on-the-fly generating a BED file with constant DP values).

## 3. Examples

Using a gold-standard 300× WGS dataset as input (details on data preparation and experimental setup are in Supplementary Information), we benchmarked samsampleX against existing tools across three scenarios: even chromosome-wide downsampling to 30×; pattern-aware downsampling of the difficult-to-sequence *HLA-A* gene to 30×; and downsampling to a realistic WEGS configuration (100× WES and 5× WGS) in the biologically important *TP53* gene. Performance was evaluated by comparing the absolute errors (absolute differences) and biases (signed differences) between the means (*μ*_*i*_) and standard deviations (*σ*_*i*_) of DP at each base pair *i* across multiple replicates. These metrics were further aggregated into mean absolute errors (MAE) and mean biases (MB) for both means and standard deviations of DP across all base pairs.

When downsampling chromosome 21 from 300× to an even 30× coverage, all tested tools showed biases in mean DP close to zero (MB(*μ*) ranging from −0.001 to 0.05, **Supplementary Table 1**), indicating that each tool was very close to the desired average target DP in downsampled data. However, the MAE in DP was lower for samsampleX, which deviated from the target by an average of only 2 reads, compared to 8 reads for the other tools (MAE(*μ*) = 2.080 vs MAE(*μ*) = 7.837 - 7.844). samsampleX also exhibited variability between downsampling replicates similar to other tools (MAE(*σ*) = 5.008 vs MAE(*σ*) = 4.876 - 4.966). These results suggest that samsampleX achieves more even coverage than uniform downsampling approaches in this scenario.

The biologically important *HLA-A* gene region is known as a difficult-to-sequence region, as reflected by a pronounced drop in coverage in a typical 30× experiment in the 1000 Genomes Project (**Supplementary Figure 1**). When simulating this region using downsampling, samsampleX showed lower bias (i.e., closer to 0) in mean DP than other tools (MB(*μ*) = 0.278 vs MB(*μ*) = −1.882 - −1.345), **Supplementary Table 2**). Although the difference in mean DP bias between samsampleX and other tools may be small, samsampleX deviated from realistic coverage by only 1 read on average, whereas the other tools deviated by 10 or more reads (MAE(￼) = 1.389 vs MAE(￼) = 10.227–10.621). Similarly to the previous experiment, samsampleX had variability between experiments comparable to other tools (MB(*σ*) = 4.122 vs MB(*σ*) = 3.668 - 4.263; MAE(*σ*) = 4.159 vs MAE(*σ*) = 3.779 – 4.301). Visual assessment of downsampling using traditional tools (**Figure 1A**) and samsampleX (**Figure 1B**) shows that traditional tools oversampled the difficult-to-sequence region while undersampling the remaining parts.

**Figure 1.**
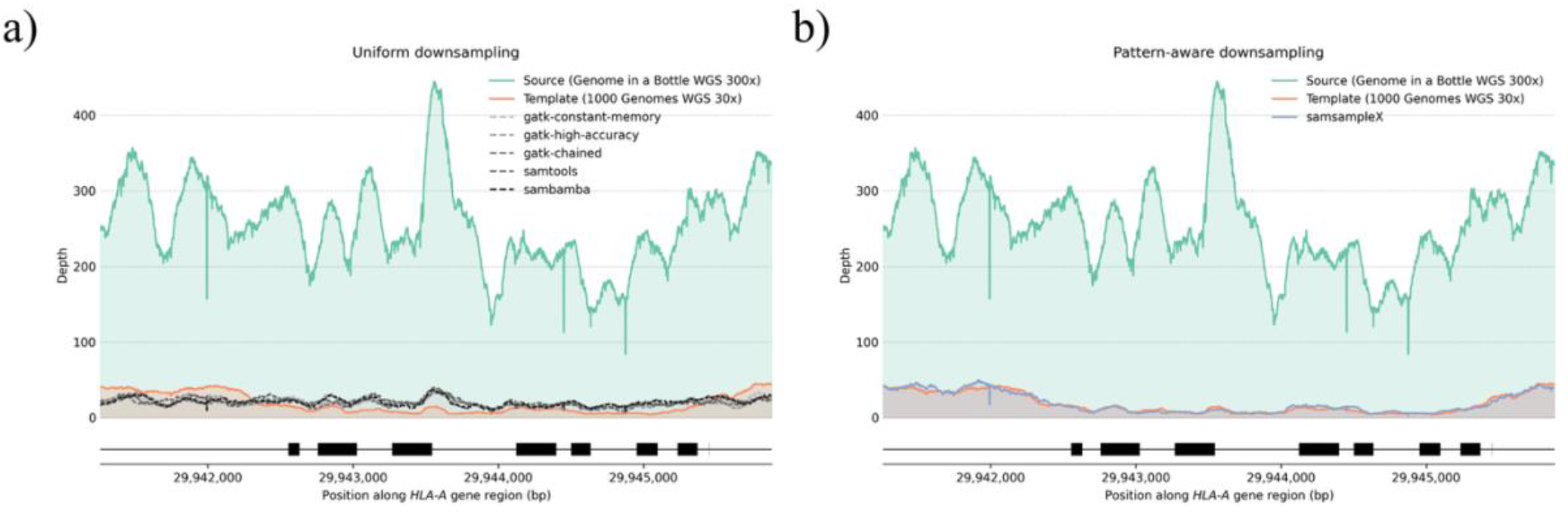
Downsampling to coverage in a typical sequencing experiment in the *HLA-A* gene region. The *HLA-A* gene region was defined as chr6:29,941,260-29,945,884 using the GRCh38 human genome build. Coding sequences in the region are annotated in black using release 49. The downsampling was performed on a Genome In A Bottle (GIAB) sample, HG002, sequenced at 300× (labeled in green as the source). The coverage representing a typical sequencing experiment was generated from 10 random samples from the 1000 Genomes Project sequenced in a 30× whole-genome experiment (labeled in orange as the template). Lines represent the mean depth computed over 10 downsampling runs with different random seeds. **A**. Uniform downsampling using traditional tools with a downsampling coefficient set to match the mean depth in the 1000 Genomes Project. **B**. A distribution-aware downsampling using samsampleX set to match a typical sequencing coverage in the 1000 Genomes Project.

samsampleX performed well when simulating realistic hybrid WEGS coverage, following high DP near coding regions and low DP in non-coding regions in the *TP53* gene (**Supplementary Figure 2**). The positive bias (MB(*μ*) = 12.601), which is higher relative to the previous scenarios, is driven by the spikes around coding regions, where samsampleX was undersampling (**Supplementary Table 3**). The variability between downsampling runs using samsampleX was similar to previous scenarios (MAE(*σ*) = 4.339). Although other tools were not designed for such a downsampling scenario, we nonetheless employed them with a sampling coefficient set to match the region-wide mean DP, in the interest of experimental consistency. We were aware, however, that their MB and MAE metrics would be significantly worse than those of samsampleX (**Supplementary Table 3**).

## 4. Runtime and memory

Our pattern-aware downsampling algorithm introduces an additional O(1) operation per read to compute the downsampling coefficient (assuming comparable read lengths) relative to uniform downsampling tools. Runtime benchmarking across the scenarios described above suggests that this added computational cost is negligible within the context of an I/O-bound workflow. Despite being implemented in Python, samsampleX achieved adequate performance characteristics suitable for large-scale applications, even relative to Java or C-based uniform downsampling tools. For example, downsampling chromosome 21 from 300× to 30× required 498 seconds and 4 GB of memory with samsampleX, and 75 - 310 seconds and <1-8 GB with other tools (**Supplementary Table 1**). The generation of BED files with target DP values for samsampleX is a separate step performed only once using the well-documented and efficient samtools tool, so it was not included in our runtime comparisons.

## 5. Conclusions

Popular bioinformatics tools with NGS downsampling functionality, such as GATK Picard, samtools, and sambamba, allow researchers to simulate NGS data across different DP ranges for benchmarking purposes. These tools typically apply uniform sampling of aligned sequencing reads, assuming equal sampling probability across the genome. While highly useful in many scenarios, this strategy may not be adequate for certain applications. In this work, we demonstrated that uniform sampling may be less accurate when simulating even DP profiles, realistic coverage patterns in hard-to-sequence regions, and hybrid NGS datasets in our tested scenarios. To address this limitation, we developed samsampleX, a novel downsampling algorithm that adjusts read sampling probabilities according to user-defined DP distribution patterns. These patterns can be derived from real-world sequencing datasets, the availability of which continues to grow, and subsequently used to generate realistic *in silico* datasets for benchmarking applications.

This work has several limitations. First, the proposed approach is designed for a specific downsampling application: reducing DP from ultra-high-DP reference datasets such as GIAB. Because such datasets are generated using exceptionally rigorous experimental design, quality control, and technical replication, their DP distribution patterns may not accurately reflect those of typical research sequencing experiments at lower DP. In cases where researchers have access to ultra-high-DP datasets generated using the same sequencing platform and experimental design as their target application, conventional uniform downsampling may be preferable, as it naturally preserves the original DP distribution. Second, we evaluated samsampleX only on short-read sequencing data, which currently remains more widely used than long-read sequencing. Although the underlying mathematical framework is applicable to long-read data in principle, additional validation is required before routine use in long-read sequencing applications.

samsampleX is a free, open-source, Python-based package that is easy to use and has computational requirements comparable to widely used tools such as GATK Picard, samtools, and sambamba. Rather than replacing these established tools, samsampleX extends existing downsampling strategies by providing additional flexibility for specialized and non-standard NGS benchmarking scenarios.

## Acknowledgements

This research was enabled in part by support provided by Calcul Québec (calculquebec.ca) and the Digital Research Alliance of Canada (alliancecan.ca).

## Data and Code Availability

samsampleX source code, supporting documentation and a benchmarking workflow are available on GitHub (github.com/sdemiriz/samsampleX) under an MIT License.

