## Supplementary Information for "samsampleX: Distribution-aware downsampling for benchmarking next-generation sequencing data"

#### 1. Per-read downsampling coefficients and their properties

Let $R$ denote the set of aligned reads and, for each genomic position $i = 1, \ldots, N$, let $R_i \subseteq R$ be the subset of reads overlapping position $i$, with observed depth $c_i = |R_i|$. Given a target depth $d_i \leq c_i$, let us define the per-position downsampling coefficient $k_i = \frac{d_i}{c_i}$.

For a read $r \in R$ of length $L_r$ spanning genomic positions $s_r, s_r + 1, s_r + 2, \ldots, s_r + L_r - 1$, let us define the per-read retention probability (i.e. per-read downsampling coefficient) as the mean of the per-position downsampling coefficients it covers

$$k_r = \frac{1}{L_r} \sum_{i=s_r}^{s_r+L_r-1} k_i$$

Each read is retained independently: a hash function maps read name to a pseudo-random value $u(r) \sim \text{Uniform}(0, 1)$, deterministic for a given read name, and the read is kept if $u(r) < k_r$. Let the random variable $X_r \sim Bernoulli(k_r)$ indicate if the read was downsampled. Then, the downsampled depth at position $i$ is

$$DP_i = \sum_{r \in R_i} X_r$$

and its expected value is

$$\mathbb{E}[DP_i] = \sum_{r \in R_i} \mathbb{E}\,[X_r] = \sum_{r \in R_i} k_r$$

When $k_i$ is constant across all genomic positions, then for any read $r \in R_i$ by definition $k_r = k_i$ and $X_r \sim Bernoulli(k_i)$ and

$$\mathbb{E}_{\text{const}}[DP_i] = \sum_{r \in R_i} k_i = c_i \cdot k_i = c_i \cdot \frac{d_i}{c_i} = d_i$$

Thus, the expected downsampled depth when $k_i$ is constant at every position exactly equals the target depth.

When $k_i$ varies across the genome, reads overlapping the same position $i$ may have different retention probabilities $k_r$, since each read averages over the genomic interval it spans. However, in short-read sequencing data, coverage profiles usually vary smoothly over distances comparable to a read length (~150-300 bp). Consequently, for reads overlapping the same genomic position, the corresponding values of $k_r$ are expected to differ only slightly. This implies that $k_r$ can be viewed as a small perturbation around the local values of $k_i$. As a result, the expected depth in the general case can be compared to the constant case by writing

$$\mathbb{E}[DP_i] - \mathbb{E}_{const}[DP_i] = \sum_{r \in R_i} k_r - \sum_{r \in R_i} k_i = \sum_{r \in R_i} (k_r - k_i)$$

Or it can also be expressed as a bias with respect to the target depth as follows

$$\mathbb{E}[DP_i] - d_i = \sum_{r \in R_i} (k_r - k_i)$$

This bias is bound by the sum of maximal differences between downsampling coefficients over positions spanned by each read

$$|\mathbb{E}[DP_i] - d_i| \leq \sum_{r \in R_i} \max_{s_r \leq j \leq s_r + L_r - 1} |k_j - k_i|$$

When $k_i$ varies slowly over the scale of read length, this bias is negligible.

**2. Data**

As an ultra-high-DP reference dataset for downsampling, we used the HG002 (NA24385) sample from Genome In A Bottle (GIAB) (Zook et al. 2016), which was sequenced to 300× at whole-genome coverage. The sequencing data were downloaded in the BAM file format from https://ftp.ncbi.nlm.nih.gov/ReferenceSamples/giab/data/AshkenazimTrio/HG002_NA24385_son/NIST_HiSeq_HG002_Homogeneity-10953946/NHGRI_Illumina300X_AJtrio_novoalign_bams/HG002.GRCh38.300x.bam.

To generate a template with DP distributions to represent typical high-DP sequencing, we used 10 randomly selected samples from the 1000 Genomes Project, which were sequenced to 30× coverage: HG01363, HG01941, HG02122, HG02127, HG02291, HG02678, HG03593, HG03780, HG03965, and NA18547. No filters were used when computing DP. At each position, DP values across all samples were averaged. The sequencing data were downloaded in the BAM file format from https://ftp.1000genomes.ebi.ac.uk/vol1/ftp/data_collections/1000G_2504_high_coverage/1000G_2504_high_coverage.sequence.index.

To generate templates with DP distributions to represent typical hybrid sequencing data, we used HG002 (NA24385) sample replicate 1, which was sequenced using WEGS 8P5× technology at

100× coverage in coding regions and 5× coverage in non-coding regions. Data is accessible at BioProject ID PRJNA1043666 on the NCBI Sequence Read Archive (SRA) (Bhérer et al. 2024).

**2. Metrics for downsampling comparison**

Downsampling with each tool was repeated 10 times using different random seeds to account for variation across downsampling runs. The DP at each position in the downsampled data was computed using samtools (Danecek et al. 2021) without any filters. For each position $i = 1, \ldots, N$ and for each tool, we computed the mean $\mu_{i,tool}$ and standard deviation $\sigma_{i,tool}$ of downsampled DP. Then, to compare results from each downsampling tool to the target, we defined the mean absolute error (MAE) and mean bias (MB) for the mean and standard deviation as

$$MAE_{tool}(\mu) = \frac{1}{N}\sum_{i=0}^{N} |\mu_{i,target} - \mu_{i,tool}|$$

$$MAE_{tool}(\sigma) = \frac{1}{N}\sum_{i=0}^{N} |\sigma_{i,target} - \sigma_{i,tool}|$$

$$MB_{tool}(\mu) = \frac{1}{N}\sum_{i=0}^{N} (\mu_{i,target} - \mu_{i,tool})$$

$$MB_{tool}(\sigma) = \frac{1}{N}\sum_{i=0}^{N} (\sigma_{i,target} - \sigma_{i,tool})$$

When comparing against the typical coverage in the 1000 Genomes Project, the $\mu_{i,target}$ and $\sigma_{i,target}$ were computed from 10 randomly selected samples. When the target was an even distribution (e.g. 30×) across the genome, then $\sigma_{i,target}$ was set to 0. When comparing to a target DP computed only using one sample (e.g. WEGS), then $\sigma_{i,target}$ was set to 0 as well.

**4. Software and hardware configuration**

All benchmarks were run on two CPU cores with 16GB of memory. Benchmarking was performed using the benchmarking functionality of Snakemake (Mölder et al. 2025) workflow manager; the full workflow is included in the accompanying GitHub repository. Additional benchmarking scenarios can be easily set up by extending the workflow configuration according to the provided documentation.

The reported runtime and memory usage metrics correspond to the CPU time elapsed and the maximum resident set size reached only by the downsampling functions from each tool. The latter metric is originally reported in megabytes (MB), converted to gigabytes (GB) for reporting.

GATK version 4.6.2.0 with HTSJDK version 4.2.0 and Picard version 3.4.0, samtools version 1.23 with htslib 1.23, and sambamba version 1.0.1 were used for performance benchmarks. samsampleX dependencies included xxhash version 3.5.0, pysam version 0.23.3, numpy version 2.3.3, and matplotlib version 3.10.8.

#### 5. Supplementary tables

Coding sequence annotations used in benchmark plot annotations are sourced from data from GENCODE (Mudge et al. 2025) release 49.

**Supplementary Table 1. Downsampling to an even 30× coverage at chromosome 21.** The downsampling was performed on a Genome In A Bottle (GIAB) sample HG002 sequenced at 300×. The downsampling coefficient for GATK-based and sambamba tools was derived from the total read counts in GIAB sample, computed using samtools (no filters). Experiments were run on two CPU cores with 16 GB of memory. Means and standard deviations (SD) were computed over 10 downsampling runs with different random seeds. MAE(μ) – mean absolute error (absolute difference) between target and downsampled mean depths. MB(μ) – mean bias (signed difference) between target and downsampled mean depths. MAE(σ) – mean absolute error (absolute difference) between the standard deviations of the target and the downsampled depths. MB(σ) – mean bias (signed difference) between standard deviations of the target and downsampled depths. In this experiment, the standard deviation of the target depth was set to 0 when computing MAE(σ) and MB(σ), because the target DP was a constant.

| Tool | Downsampling coefficient | Target depth | Mean runtime [s] (SD) | Mean memory [GB] (SD) | Mean downsampled depth (SD) | MAE(μ) | MAE(σ) | MB(μ) | MB(σ) |
|---|---|---|---|---|---|---|---|---|---|
| **GATK Constant Memory** | 0.094 | - | 127.629 (01.244) | 0.931 (0.003) | 29.998 (4.959) | 7.844 | 4.959 | 0.002 | -4.959 |
| **GATK High Accuracy** | 0.094 | - | 309.244 (07.574) | 8.456 (0.077) | 30.001 (4.876) | 7.839 | 4.876 | -0.001 | -4.876 |
| **GATK Chained** | 0.094 | - | 158.882 (10.355) | 3.435 (0.105) | 30.000 (4.951) | 7.843 | 4.951 | 0.000 | -4.951 |
| **samtools** | 0.094 | - | 74.895 (00.387) | 0.047 (0.001) | 29.997 (4.962) | 7.843 | 4.962 | 0.003 | -4.962 |
| **sambamba** | 0.094 | - | 159.647 (25.372) | 0.048 (0.002) | 29.997 (4.966) | 7.837 | 4.966 | 0.003 | -4.966 |
| **samsampleX** | - | 30.000 | 498.179 (28.157) | 4.108 (1.475) | 29.950 (5.008) | 2.080 | 5.008 | 0.050 | -5.008 |

**Supplementary Table 2. Downsampling to coverage in a typical sequencing experiment in the *HLA-A* gene region.** The downsampling was performed on a Genome In A Bottle (GIAB) sample, HG002, sequenced at 300×. The *HLA-A* gene region was defined as chr6:29,941,260-29,945,884 region using the GRCh38 human genome build version. The coverage template for samsampleX representing a typical sequencing experiment was generated using 10 random samples from the 1000 Genomes Project sequenced in 30× whole-genome sequencing experiment. The downsampling coefficient for GATK-based and sambamba tools was derived from the average total read counts in input dataset (i.e., 1000 Genomes Project) computed using samtools (no filters). Downsampling experiments were run on two CPU cores with 16 GB of memory. Means and standard deviations (SD) were computed over 10 runs with different random seeds. MAE(μ) – mean absolute error (absolute difference) between target and downsampled mean depths. MB(μ) – mean bias (signed difference) between target and downsampled mean depths. MAE(σ) – mean absolute error (absolute difference) between the standard deviations of the target and the downsampled depths. MB(σ) – mean bias (signed difference) between standard deviations of the target and downsampled depths.

| Tool | Downsampling coefficient | Mean template depth (SD) | Mean runtime [s] (SD) | Mean memory [GB] (SD) | Mean downsampled depth (SD) | MAE(μ) | MAE(σ) | MB(μ) | MB(σ) |
|---|---|---|---|---|---|---|---|---|---|
| **GATK Constant Memory** | 0.078 | - | 4.368 (0.345) | 0.295 (0.004) | 20.159 (4.209) | 10.334 | 3.779 | -1.345 | 3.668) |
| **GATK High Accuracy** | 0.078 | - | 4.424 (0.196) | 0.296 (0.006) | 20.112 (3.998) | 10.621 | 4.015 | -1.299 | 3.879 |
| **GATK Chained** | 0.078 | - | 4.167 (0.242) | 0.296 (0.003) | 19.786 (3.614) | 10.227 | 4.301 | -0.972 | 4.263 |
| **samtools** | 0.078 | - | 0.183 (0.090) | 0.037 (0.012) | 19.695 (4.014) | 10.237 | 3.921 | -0.882 | 3.862 |
| **sambamba** | 0.078 | - | 0.058 (0.063) | 0.032 (0.013) | 20.034 (4.003) | 10.255 | 3.921 | -1.220 | 3.874 |
| **samsampleX** | - | 18.813 (7.877) | 0.529 (0.112) | 0.074 (0.008) | 18.536 (3.755) | 1.389 | 4.159 | 0.278 | 4.122 |

**Supplementary Table 3. Downsampling to coverage in a hybrid sequencing experiment in the *TP53* gene region.** The downsampling was performed on a Genome In A Bottle (GIAB) sample, HG002, sequenced at 300×. The *TP53* gene region was defined as chr17:7758460-7784220 region using the GRCh38 human genome build version. The coverage template for samsampleX representing a hybrid sequencing experiment was created from a sample sequenced using the Whole Genome Exome Sequencing (WEGS) approach, with 100× coverage in coding sequences and 5× coverage in non-coding sequences. The downsampling coefficient for GATK-based and sambamba tools was derived from the total read counts in input dataset (i.e., WEGS), computed using samtools (no filters). Means and standard deviations (SD) were computed over 10 downsampling runs with different random seeds. MAE(μ) – mean absolute error (absolute difference) between target and downsampled mean depths. MB(μ) – mean bias (signed difference) between target and downsampled mean depths. MAE(σ) – mean absolute error (absolute difference) between the standard deviations of the target and the downsampled depths. MB(σ) – mean bias (signed difference) between standard deviations of the target and downsampled depths. In this experiment, the standard deviation of the target depth was set to 0 when computing MAE(σ) and MB(σ), because the target DP was generated using single sample.

| Tool | Downsampling coefficient | Template depth | Mean runtime [s] (SD) | Mean memory [GB] (SD) | Mean downsampled depth (SD) | MAE(μ) | MAE(σ) | MB(μ) | MB(σ) |
|---|---|---|---|---|---|---|---|---|---|
| **GATK Constant Memory** | 0.289 | - | 4.721 (0.558) | 0.310 (0.017) | 75.840 ( 7.241) | 60.870 | 7.241 | -30.360 | -7.241 |
| **GATK High Accuracy** | 0.289 | - | 4.992 (0.234) | 0.327 (0.024) | 76.032 ( 7.133) | 61.091 | 7.133 | -30.552 | -7.133 |
| **GATK Chained** | 0.289 | - | 4.574 (0.548) | 0.304 (0.015) | 75.970 ( 7.294) | 60.952 | 7.294 | -30.490 | -7.294 |
| **samtools** | 0.289 | - | 0.268 (0.071) | 0.042 (0.004) | 76.521 ( 7.157) | 61.379 | 7.157 | -31.040 | -7.157 |
| **sambamba** | 0.289 | - | 0.229 (0.127) | 0.040 (0.001) | 76.169 ( 7.384) | 61.140 | 7.384 | -30.689 | - 7.384 |
| **samsampleX** | - | 45.852 | 0.736 (0.098) | 0.079 (0.002) | 32.880 ( 4.181) | 23.960 | 4.181 | 12.601 | - 4.181 |

### 6. Supplementary figures

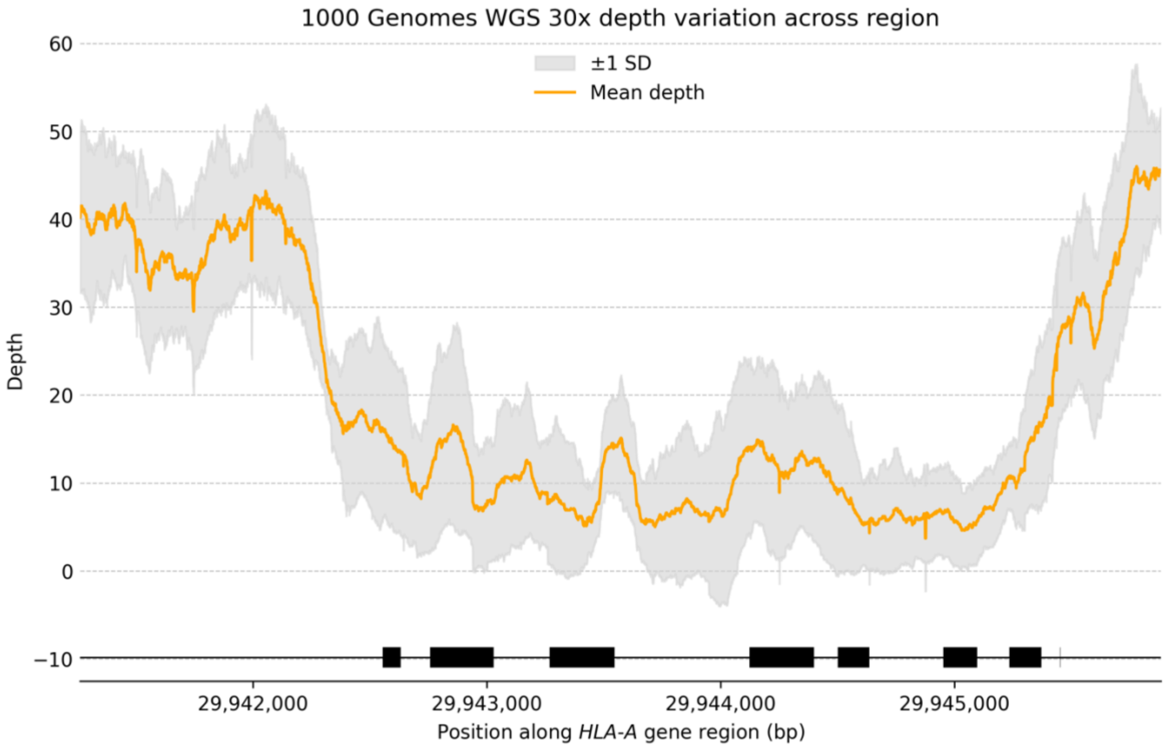


**Supplementary Figure 1. Coverage in a typical sequencing experiment in the *HLA-A* gene region.** The *HLA-A* gene region was defined as chr6:29,941,260-29,945,884 region using the GRCh38 human genome build version. Coding sequences in the region are annotated in black using GENCODE (Mudge et al. 2025) release 49 on human genome build GRCh38. The coverage representing a typical sequencing experiment was generated using 10 random samples from the 1000 Genomes Project sequenced in 30× whole-genome sequencing experiment. SD - standard deviation.

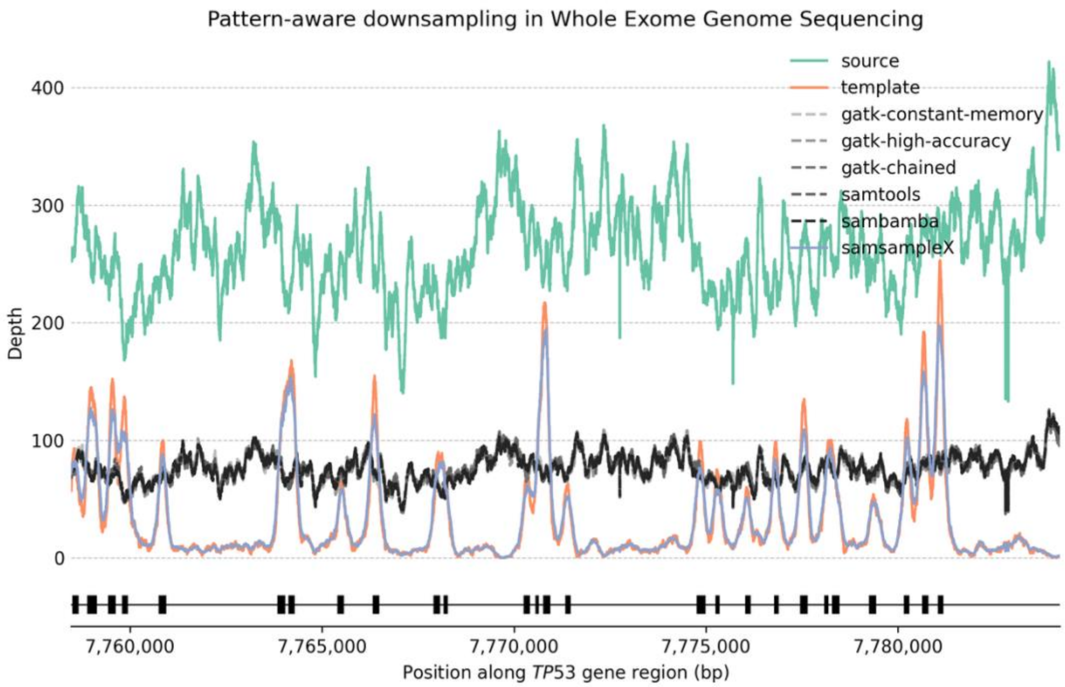


**Supplementary Figure 2. Downsampling to coverage in a hybrid sequencing experiment in the *TP53* gene region.** The *TP53* gene region was defined as chr17:7758460-7784220 region using the GRCh38 human genome build version. Coding sequences in the region are annotated in black using GENCODE (Mudge et al. 2025) release 49 on human genome build GRCh38. The downsampling was performed on a Genome In A Bottle (GIAB) sample, HG002, sequenced at 300× (labeled in green as the source). The targeted coverage representing a hybrid sequencing experiment was created from a sample sequenced using the Whole Genome Exome Sequencing (WEGS) approach, with 100× coverage in coding sequences and 5× coverage in non-coding sequences (labeled in orange as the template). The downsampling coefficient for GATK-based and sambamba tools was derived from the total read counts in the input dataset (i.e., WEGS), computed using samtools (no filters). Lines represent mean depth computed over 10 downsampling runs with different random seeds.
